## Supplementary for "Gradual Compaction of the Nascent Peptide During Cotranslational Folding on the Ribosome"

### Supplementary Tables

**Supplementary Table 1** ACF (each average of N≥8) fits of HemK constructs in solution. All fit errors are calculated as standard error of the mean and are <10 %. Rates $k_1$ and $k_d$ are in $s^{-1}$. $\tau_1$ and $\tau_{d1}$ are relaxation time constants of the respective exponents, in s, $\tau$=1/k. N is the average number of molecules in the confocal volume and $c_1$ is the amplitude of the fast relaxation time.

| HemK | $c_1$ | $\tau_1$ x$10^{-7}$ | $k_1$ x$10^6$ | N | $\tau_{d1}$ x$10^{-5}$ | $k_d$ x$10^3$ |
|---|---|---|---|---|---|---|
| 0% glycerol | | | | | | |
| 70 W6 | | | | 0.83 | 5.16 | 19.38 |
| 70 W6F | | | | 0.83 | 5.42 | 18.43 |
| 70 4xA W6 | | | | 0.85 | 5.78 | 17.30 |
| 70 4xA W6F | | | | 0.83 | 5.59 | 17.89 |
| 14 W6 | | | | 0.84 | 5.88 | 17.00 |
| 14 W6F | | | | 0.86 | 6.61 | 15.13 |
| 50% glycerol | | | | | | |
| 70 W6 | | | | 0.94 | 30.78 | 3.25 |
| 70 W6F | | | | 0.95 | 33.87 | 2.95 |
| 70 4xA W6 | | | | 0.95 | 30.40 | 3.29 |
| 70 4xA W6F | | | | 0.95 | 29.10 | 3.44 |
| 14 W6 | 0.15 | 5.13 | 1.95 | 0.97 | 45.09 | 2.22 |
| 14 W6F | | | | 0.97 | 60.31 | 1.66 |

**Supplementary Table 2** Results of analytical fits of PET-FCS ACF (each ACF an average of N≥8) for RNCs. All fit errors are calculated as standard error of the mean and are indicated in the table. Rates ($k_x$) are in $s^{-1}$. $\tau_1$, $\tau_2$, $\tau_f$ and $\tau_d$ are relaxation time constants of the respective exponents, in s, $\tau$=1/k.

| Construct | $c_1$ | $k_1$ x$10^6$ | $\tau_1$ x$10^{-7}$ | $c_2$ | $k_2$ x$10^5$ | $\tau_2$ x$10^{-6}$ | F | $k_f$ x$10^4$ | $\tau_f$ x$10^{-5}$ | N | $k_d$ x$10^3$ | $\tau_d$ |
|---|---|---|---|---|---|---|---|---|---|---|---|---|
| 70 W6 | 0.17 ± 0.01 | 2.58 ± 0.21 | 3.87 | 0.68 ± 0.01 | 4.55 ± 0.09 | 2.20 | 0.18 ± 0.003 | 2.29 ± 0.09 | 4.38 | 0.93 ± 0.003 | 1.05 ± 0.008 | 0.001 |
| 70 W6F | 0.14 ± 0.01 | 2.29 ± 0.19 | 4.36 | 0.62 ± 0.01 | 4.59 ± 0.09 | 2.18 | 0.13 ± 0.002 | 2.45 ± 0.10 | 4.09 | 0.93 ± 0.002 | 0.95 ± 0.005 | 0.001 |
| 70 4xA W6 | 0.18 ± 0.01 | 2.67 ± 0.16 | 3.74 | 0.69 ± 0.01 | 4.51 ± 0.07 | 2.22 | 0.16 ± 0.002 | 2.45 ± 0.09 | 4.08 | 0.93 ± 0.002 | 0.99 ± 0.006 | 0.001 |
| 70 4xA W6F | 0.13 ± 0.01 | 2.61 ± 0.23 | 3.84 | 0.61 ± 0.01 | 4.68 ± 0.09 | 2.13 | 0.12 ± 0.002 | 2.86 ± 0.14 | 3.49 | 0.92 ± 0.002 | 0.93 ± 0.005 | 0.001 |
| 102 W6 | 0.20 ± 0.01 | 3.30 ± 0.19 | 3.03 | 0.61 ± 0.01 | 4.29 ± 0.07 | 2.33 | 0.18 ± 0.002 | 2.36 ± 0.09 | 4.23 | 0.94 ± 0.002 | 0.95 ± 0.007 | 0.001 |
| 102 W6F | 0.14 ± 0.01 | 2.91 ± 0.23 | 3.43 | 0.56 ± 0.01 | 4.66 ± 0.09 | 2.15 | 0.13 ± 0.002 | 2.54 ± 0.11 | 3.94 | 0.93 ± 0.002 | 0.93 ± 0.005 | 0.001 |
| 102 loop | 0.13 ± 0.01 | 2.89 ± 0.24 | 3.46 | 0.57 ± 0.01 | 4.33 ± 0.08 | 2.31 | 0.14 ± 0.002 | 2.54 ± 0.12 | 3.93 | 0.93 ± 0.002 | 0.86 ± 0.005 | 0.001 |
| 112 loop | 0.14 ± 0.01 | 2.72 ± 0.21 | 3.68 | 0.54 ± 0.01 | 4.24 ± 0.08 | 2.36 | 0.13 ± 0.002 | 2.47 ± 0.11 | 4.05 | 0.93 ± 0.002 | 0.88 ± 0.005 | 0.001 |
| 112 W6 | 0.19 ± 0.01 | 3.61 ± 0.19 | 2.77 | 0.48 ± 0.01 | 4.41 ± 0.08 | 2.27 | 0.10 ± 0.002 | 2.79 ± 0.16 | 3.59 | 0.93 ± 0.002 | 0.84 ± 0.004 | 0.001 |
| 112 W6F | 0.14 ± 0.01 | 2.94 ± 0.22 | 3.40 | 0.52 ± 0.01 | 4.45 ± 0.09 | 2.25 | 0.11 ± 0.002 | 2.51 ± 0.13 | 3.98 | 0.93 ± 0.002 | 0.91 ± 0.005 | 0.001 |
| 112 4xA W6 | 0.20 ± 0.01 | 2.16 ± 0.13 | 4.63 | 0.59 ± 0.01 | 3.73 ± 0.08 | 2.68 | 0.21 ± 0.003 | 1.76 ± 0.06 | 5.68 | 0.95 ± 0.003 | 1.00 ± 0.008 | 0.001 |
| 112 4xA W6F | 0.12 ± 0.01 | 2.52 ± 0.23 | 3.97 | 0.43 ± 0.01 | 3.94 ± 0.97 | 2.54 | 0.11 ± 0.002 | 2.30 ± 0.13 | 4.35 | 0.93 ± 0.002 | 0.86 ± 0.005 | 0.001 |

**Supplementary Table 3** Results of global fitting of the free Trp titration (dataset A) to model 5e-H. Rates that were linked during global fit are shown in the same cell shades. All reported rates (k) are in $\mu s^{-1}$, except the Wd and Wc k on rates that are in $mM^{-1}$ $\mu s^{-1}$, rates where SEM exceeds value are given as not significant (n.s.). A covariance matrix derived using nonlinear regression algorithms is used to estimate the standard errors (SEM) by the Kintek Explorer software.

| Construct | k | D ↔ Rd | C ↔ Rc | D ↔ Wd | C ↔ Wc | D ↔ C |
|---|---|---|---|---|---|---|
| 70 W6F | on | 139 ± 14 | n.s. | 0.010 ± 0.004 | ~ 0 | n.s. |
| | off | n.s. | n.s. | 2.2 ± 0.2 | 2.2 ± 0.2 | 0.50 ± 0.07 |
| 102 W6F | on | 461 ± 69 | n.s. | 0.03 ± 0.01 | ~ 0 | n.s. |
| | off | n.s. | 0.51 ± 0.09 | 2.2 ± 0.2 | 2.2 ± 0.2 | 1.9 ± 0.2 |

**Supplementary Table 4** Results of global fitting of the free Trp titration (dataset A) to model 5e-O. Legend as in Supplementary Table 3.

| Construct | k | D ↔ R | C ↔ R | D ↔ W | C ↔ W | D ↔ C |
|---|---|---|---|---|---|---|
| 70 W6F | on | 83 ± 17 | 0.5 ± 0.06 | 0.006 ± 0.003 | ~ 0 | n.s. |
| | off | n.s. | n.s. | 2 ± 0.1 | 2 ± 0.1 | n.s. |
| 102 W6F | on | 85 ± 15 | 0.5 ± 0.05 | 0.007 ± 0.003 | ~ 0 | n.s. |
| | off | n.s. | n.s. | 2 ± 0.1 | 2 ± 0.1 | n.s. |

**Supplementary Table 5** Results of global fitting of the dataset B to the model 5e-H. Rates (k) are reported in μs$^{-1}$, rates linked during global fit are shown in the same cell shade; locked values are in red. A covariance matrix derived using nonlinear regression algorithms is used to estimate the standard errors (SEM) by the Kintek Explorer software.

| Construct | k | D ↔ R | C ↔ R | D ↔ W | C ↔ W | D ↔ C |
|---|---|---|---|---|---|---|
| 70 wt | on | 9.7 ± 1.3 | 0.24 ± 0.02 | 7.1 ± 1.2 | 0.04 ± 0.02 | 1.4 ± 0.1 |
| | off | 0.30 ± 0.02 | 0.30 ± 0.02 | 2.2 | 2.2 | 0.18 ± 0.01 |
| 70 4xA | on | 9.7 ± 1.3 | 0.24 ± 0.02 | 7.1 ± 1.2 | 0.04 ± 0.02 | 1.4 ± 0.1 |
| | off | 0.30 ± 0.02 | 0.30 ± 0.02 | 2.2 | 2.2 | 0.18 ± 0.01 |
| 102 wt | on | 4.7 ± 0.2 | 0.24 ± 0.02 | 3.4 ± 0.2 | 0.04 ± 0.02 | 0.50 ± 0.04 |
| | off | 0.30 ± 0.02 | 0.30 ± 0.02 | 2.2 | 2.2 | 0.14 ± 0.01 |
| 102 loop | on | 4.7 ± 0.2 | 0.24 ± 0.02 | | | 5.04 ± 0.73 |
| | off | 0.30 ± 0.02 | 0.30 ± 0.02 | | | 1.4 ± 0.2 |
| 112 loop | on | 11.5 ± 0.5 | 0.24 ± 0.02 | | | 3.7 ± 0.4 |
| | off | 0.30 ± 0.02 | 0.30 ± 0.02 | | | 0.41 ± 0.03 |
| 112 wt | on | 11.5 ± 0.5 | 0.24 ± 0.02 | 8.4 ± 0.4 | 0.04 ± 0.02 | 0.04 ± 0.11 |
| | off | 0.30 ± 0.02 | 0.30 ± 0.02 | 2.2 | 2.2 | 0.005 ± 0.018 |
| 112 4xA | on | 11 ± 2.9 | 0.22 ± 0.08 | 8.1 ± 2.5 | 0.04 ± 0.07 | 2.2 ± 0.1 |
| | off | 0.30 ± 0.02 | 0.30 ± 0.02 | 2.2 | 2.2 | 0.23 ± 0.05 |

**Supplementary Table 6** Results of global fitting of the dataset B to the model 5e-O. The legend is the same as in Supplementary Table 5.

| Construct | k | D ↔ R | C ↔ R | D ↔ W | C ↔ W | D ↔ C |
|---|---|---|---|---|---|---|
| 70 wt | on | 22 ± 3.3 | 0.37 ± 0.02 | 4.2 ± 3 | 0.07 ± 0.03 | 0.975 ± 0.3 |
| | off | 0.3 ± 0.02 | 0.3 ± 0.02 | 2 | 2 | 0.0167 ± 0.02 |
| 70 4xA | on | 22 ± 3.3 | 0.37 ± 0.02 | 4.2 ± 3 | 0.07 ± 0.03 | 0.975 ± 0.3 |
| | off | 0.3 ± 0.02 | 0.3 ± 0.02 | 2 | 2 | 0.0167 ± 0.02 |
| 102 wt | on | 25 ± 1.2 | 0.37 ± 0.02 | 4.8 ± 1 | 0.07 ± 0.03 | 0.13 ± 0.3 |
| | off | 0.3 ± 0.02 | 0.3 ± 0.02 | 2 | 2 | 0.002 ± 0.02 |
| 102 loop | on | 25 ± 1.2 | 0.37 ± 0.02 | | | 4.39 ± 0.5 |
| | off | 0.3 ± 0.02 | 0.3 ± 0.02 | | | 0.0653 ± 0.03 |
| 112 loop | on | 29 ± 1.3 | 0.37 ± 0.02 | | | 3.44 ± 0.5 |
| | off | 0.3 ± 0.02 | 0.3 ± 0.02 | | | 0.0442 ± 0.03 |
| 112 wt | on | 29 ± 1.3 | 0.37 ± 0.02 | 5.6 ± 1 | 0.07 ± 0.03 | 0.0002 ± 0.3 |
| | off | 0.3 ± 0.02 | 0.3 ± 0.02 | 2 | 2 | 0.000002 ± 0.02 |
| 112 4xA | on | 24 ± 8.2 | 0.35 ± 0.02 | 4.7 ± 8 | 0.07 ± 0.01 | 1.62 ± 0.4 |
| | off | 0.3 ± 0.02 | 0.3 ± 0.02 | 2 | 2 | 0.0232 ± 0.1 |

**Supplementary Table 7** Upper and lower boundaries of each rate parameter computed with a $\chi^2/_{min}\chi^2$ threshold: 0.8333. Rates linked during global fit are shown in the same cell shade; locked values are in red.

| Construct | Model 5e-O | | | Elemental Rate | Model 5e-H | | |
|---|---|---|---|---|---|---|---|
| | Best-fit Value | Lower Boundary | Upper Boundary | | Best-fit Value | Lower Boundary | Upper Boundary |
| 70wt | 21.8 | 17.4 | 28.9 | $k_{on}$ D ↔ $R_{(d)}$ | 9.7 | 6.6 | 16.3 |
| | 0.37 | 0.35 | 0.39 | $k_{on}$ C ↔ $R_{(c)}$ | 0.24 | 0.15 | 0.30 |
| | 0.31 | 0.25 | 0.46 | $k_{off}$ D/C ↔ $R_{(d/c)}$ | 0.30 | 0.17 | 0.45 |
| | 4.2 | 2.6 | 6.3 | $k_{on}$ D ↔ $W_{(d)}$ | 7.1 | 5.4 | 9.4 |
| | 0.07 | 0.04 | 0.10 | $k_{on}$ C ↔ $W_{(c)}$ | 0.04 | 0.001 | 0.13 |
| | 2 | n/a | n/a | $k_{off}$ D/C ↔ $W_{(d/c)}$ | 2.2 | n/a | n/a |
| | 0.98 | 0.45 | 2.83 | $k_{on}$ D ↔ C | 1.4 | 0.56 | 2.8 |
| | 0.02 | 0.01 | 0.04 | $k_{off}$ D ↔ C | 0.18 | 0.07 | 0.35 |
| 70 4xA | 21.8 | 17.4 | 28.9 | $k_{on}$ D ↔ $R_{(d)}$ | 9.7 | 6.6 | 16.3 |
| | 0.37 | 0.35 | 0.39 | $k_{on}$ C ↔ $R_{(c)}$ | 0.24 | 0.15 | 0.30 |
| | 0.31 | 0.25 | 0.46 | $k_{off}$ D/C ↔ $R_{(d/c)}$ | 0.30 | 0.17 | 0.45 |
| | 4.2 | 2.6 | 6.3 | $k_{on}$ D ↔ $W_{(d)}$ | 7.1 | 5.4 | 9.4 |
| | 0.07 | 0.04 | 0.10 | $k_{on}$ C ↔ $W_{(c)}$ | 0.04 | 0.001 | 0.13 |
| | 2 | n/a | n/a | $k_{off}$ D/C ↔ $W_{(d/c)}$ | 2.2 | n/a | n/a |
| | 0.98 | 0.45 | 2.83 | $k_{on}$ D ↔ C | 1.4 | 0.56 | 2.8 |
| | 0.02 | 0.01 | 0.04 | $k_{off}$ D ↔ C | 0.18 | 0.07 | 0.35 |
| 102 wt | 25 | 20 | 35.2 | $k_{on}$ D ↔ $R_{(d)}$ | 4.7 | 3.0 | 6.7 |
| | 0.37 | 0.35 | 0.39 | $k_{on}$ C ↔ $R_{(c)}$ | 0.24 | 0.15 | 0.30 |
| | 0.31 | 0.25 | 0.46 | $k_{off}$ D/C ↔ $R_{(d/c)}$ | 0.30 | 0.17 | 0.45 |
| | 4.9 | 2.97 | 7.2 | $k_{on}$ D ↔ $W_{(d)}$ | 3.4 | 2.7 | 4.1 |
| | 0.07 | 0.04 | 0.10 | $k_{on}$ C ↔ $W_{(c)}$ | 0.04 | 0.001 | 0.13 |
| | 2 | n/a | n/a | $k_{off}$ D/C ↔ $W_{(d/c)}$ | 2.2 | n/a | n/a |
| | 0.13 | $8x10^{-7}$ | 1.8 | $k_{on}$ D ↔ C | 0.50 | 0.08 | 0.88 |
| | 0.002 | $1x10^{-8}$ | 0.025 | $k_{off}$ D ↔ C | 0.14 | 0.03 | 0.24 |
| 102 loop | 25 | 20 | 35.2 | $k_{on}$ D ↔ $R_{(d)}$ | 4.7 | 3.0 | 6.7 |
| | 0.37 | 0.35 | 0.39 | $k_{on}$ C ↔ $R_{(c)}$ | 0.24 | 0.15 | 0.30 |
| | 0.31 | 0.25 | 0.46 | $k_{off}$ D/C ↔ $R_{(d/c)}$ | 0.30 | 0.17 | 0.45 |
| | 4.4 | 3.5 | 7.7 | $k_{on}$ (D ↔ C) | 5.04 | 1.8 | 69.8 |
| | 0.065 | 0.056 | 0.089 | $k_{off}$ (D ↔ C) | 1.4 | 0.54 | 21.1 |

| | | | | | | | |
|---|---|---|---|---|---|---|---|
| 112 loop | 29 | 23.2 | 40.7 | $k_{on}$ D ↔ $R_{(d)}$ | 11.5 | 7.8 | 20.2 |
| | 0.37 | 0.35 | 0.39 | $k_{on}$ C ↔ $R_{(c)}$ | 0.24 | 0.15 | 0.30 |
| | 0.31 | 0.25 | 0.46 | $k_{off}$ D/C ↔ $R_{(d/c)}$ | 0.30 | 0.17 | 0.45 |
| | 3.4 | 2.8 | 6.0 | $k_{on}$ D ↔ C | 3.7 | 0.91 | 8.1 |
| | 0.044 | 0.038 | 0.066 | $k_{off}$ D ↔ C | 0.41 | 0.11 | 0.71 |
| 112 wt | 29 | 23.2 | 40.7 | $k_{on}$ D ↔ $R_{(d)}$ | 11.5 | 7.8 | 20.2 |
| | 0.37 | 0.35 | 0.39 | $k_{on}$ C ↔ $R_{(c)}$ | 0.24 | 0.15 | 0.30 |
| | 0.31 | 0.25 | 0.46 | $k_{off}$ D/C ↔ $R_{(d/c)}$ | 0.30 | 0.17 | 0.45 |
| | 5.6 | 3.4 | 8.8 | $k_{on}$ D ↔ $W_{(d)}$ | 8.4 | 6.35 | 11.8 |
| | 0.07 | 0.04 | 0.10 | $k_{on}$ C ↔ $W_{(c)}$ | 0.04 | 0.001 | 0.13 |
| | 2 | n/a | n/a | $k_{off}$ D/C ↔ $W_{(d/c)}$ | 2.2 | n/a | n/a |
| | 0.0002 | $2x10^{-8}$ | 1.6 | $k_{on}$ D ↔ C | 0.04 | $5x10^{-5}$ | 0.66 |
| | $2x10^{-6}$ | $2x10^{-10}$ | 0.02 | $k_{off}$ D ↔ C | 0.005 | $6x10^{-6}$ | 0.06 |
| 112 4xA | 24.1 | 17.4 | 40 | $k_{on}$ D ↔ $R_{(d)}$ | 11 | 7.5 | 19.4 |
| | 0.35 | 0.33 | 0.36 | $k_{on}$ C ↔ $R_{(c)}$ | 0.22 | 0.14 | 0.28 |
| | 0.31 | 0.25 | 0.46 | $k_{off}$ D/C ↔ $R_{(d/c)}$ | 0.30 | 0.17 | 0.45 |
| | 4.7 | 2.9 | 8.3 | $k_{on}$ D ↔ $W_{(d)}$ | 8.1 | 6.1 | 10.1 |
| | 0.067 | 0.041 | 0.095 | $k_{on}$ C ↔ $W_{(c)}$ | 0.04 | 0.001 | 0.12 |
| | 2 | n/a | n/a | $k_{off}$ D/C ↔ $W_{(d/c)}$ | 2.2 | n/a | n/a |
| | 1.6 | 0.66 | 4.7 | $k_{on}$ D ↔ C | 2.2 | 1.6 | 3.4 |
| | 0.023 | 0.012 | 0.045 | $k_{off}$ D ↔ C | 0.2 | 0.18 | 0.34 |

**Supplementary Table 8** Free energy calculations for all RNC constructs using rates derived from the 5e-H model. Elemental rate SEM from the kinetic fits are propagated through equation 2 (Methods).

| Construct | $\Delta G°_D – \Delta G^‡$, kJ mol$^{-1}$ | $\Delta G°_C – \Delta G^‡$, kJ mol$^{-1}$ | $\Delta G°_D – \Delta G°_C$, kJ mol$^{-1}$ |
|---|---|---|---|
| 70 wt | 37.5 ± 0.0001 | 42.5 ± 0.0001 | 5.0 ± 0.0001 |
| 70 4xA | 37.5 ± 0.0001 | 42.5 ± 0.0001 | 5.0 ± 0.0001 |
| 102 wt | 40.0 ± 0.0001 | 43.2 ± 0.0001 | 3.1 ± 0.0001 |
| 102 loop | 34.4 ± 0.0001 | 37.5 ± 0.0001 | 3.1 ± 0.0001 |
| 112 loop | 35.1 ± 0.0001 | 40.5 ± 0.0001 | 5.4 ± 0.0001 |
| 112 wt | 46.2 ± 0.003 | 51.3 ± 0.004 | 5.1 ± 0.005 |
| 112 4xA | 36.4 ± 0.00005 | 42.0 ± 0.0002 | 5.5 ± 0.0002 |

**Supplementary Table 9** Free energy calculations for all RNC constructs from the 5e-O model rates. Elemental rate SEM from the kinetic fits are propagated through equation 2 (Methods).

| Construct | $\Delta G°_D – \Delta G^‡$, kJ mol$^{-1}$ | $\Delta G°_C – \Delta G^‡$, kJ mol$^{-1}$ | $\Delta G°_D – \Delta G°_C$, kJ mol$^{-1}$ |
|---|---|---|---|
| 70 wt | 38.4 ± 0.0003 | 48.4 ± 0.001 | 10.0 ± 0.001 |
| 70 4xA | 38.4 ± 0.0003 | 48.4 ± 0.0000 | 10.0 ± 0.0003 |
| 102 wt | 43.3 ± 0.002 | 53.6 ± 0.01 | 10.3 ± 0.01 |
| 102 loop | 34.7 ± 0.0001 | 45.0 ± 0.0005 | 10.3 ± 0.0005 |
| 112 loop | 35.3 ± 0.0001 | 46.0 ± 0.0007 | 10.7 ± 0.0007 |
| 112 wt | 59.2 ± 1.5 | 70.5 ± 10 | 11.3 ± 10.1 |
| 112 4xA | 37.2 ± 0.0002 | 47.6 ± 0.004 | 10.4 ± 0.004 |

**Supplementary Table 10** PET-FCS constructs aa sequences N- to C-terminus. Spacing every 10 aa.

| HemK | N- to C-terminus 112 aa constructs |
|---|---|
| **wt** | MEFQHWLREA ISQLQASESP RRDAEILLEH VTGKGRTFIL AFGETQLTDE QCQQLDALLT RRRDGEPIAH LTGVREFFSL PLFVSPATLI PRPDTECLVE QALARLPEQP CR |
| **wt W6F** | MEFQHFLREA ISQLQASESP RRDAEILLEH VTGKGRTFIL AFGETQLTDE QCQQLDALLT RRRDGEPIAH LTGVREFFSL PLFVSPATLI PRPDTECLVE QALARLPEQP CR |
| **looped** | MEFQHFLREA ISQLQASESP RRDAEILLEH VTGKGRTFIL AFGGGGGGET QLTDEQCQQL DALLTRRRDG EPIAHLTGVR EFFSLPLFVS PATLIPRPDT ECLVEQALAR LPEQPCR |
| **4xA** | MEFQHWLREA ISQLQASESP RRDAEIAAEH VTGKGRTFIL AFGETQLTDE QCQQADAALT RRRDGEPIAH LTGVREFFSL PLFVSPATLI PRPDTECLVE QALARLPEQP CR |
| **4xA W6F** | MEFQHFLREA ISQLQASESP RRDAEIAAEH VTGKGRTFIL AFGETQLTDE QCQQADAALT RRRDGEPIAH LTGVREFFSL PLFVSPATLI PRPDTECLVE QALARLPEQP CR |

**Supplementary Table 11** Force profile construct of wt HemK full-length, aa sequence N- to C-terminus; numbers indicate construct truncations in HemK aa.

| **HemK** | **SecM** | **CspA** |
|---|---|---|
| MEYQHWLREA ISQLQASESP RR$^{22}$ DA$^{24}$ EI$^{26}$ LL$^{28}$ EH$^{30}$ VT$^{32}$ GK$^{34}$ GR$^{36}$ TF$^{38}$ IL$^{40}$ AF$^{42}$ GE$^{44}$ TQ$^{46}$ LT$^{48}$ DE$^{50}$ QC$^{52}$ QQ$^{54}$ LD$^{56}$ AL$^{58}$ LT RR$^{62}$ RD$^{64}$ GE$^{66}$ PI$^{68}$ AH L$^{71}$ TG$^{73}$ VR$^{75}$ E$^{76}$ F$^{77}$ W$^{78}$ S$^{79}$ L$^{80}$ P$^{81}$ L$^{82}$ F$^{83}$ V$^{84}$ S$^{85}$ P$^{86}$ A$^{87}$ T$^{88}$ L$^{89}$ I$^{90}$ P$^{91}$ RP$^{93}$ DT$^{95}$ EC$^{97}$ LV$^{99}$ E Q$^{101}$ | FSTPVWIS QAQGIRAGP | MSGKMTGIVK WFNADKGFGFITP |

### Supplementary Figures

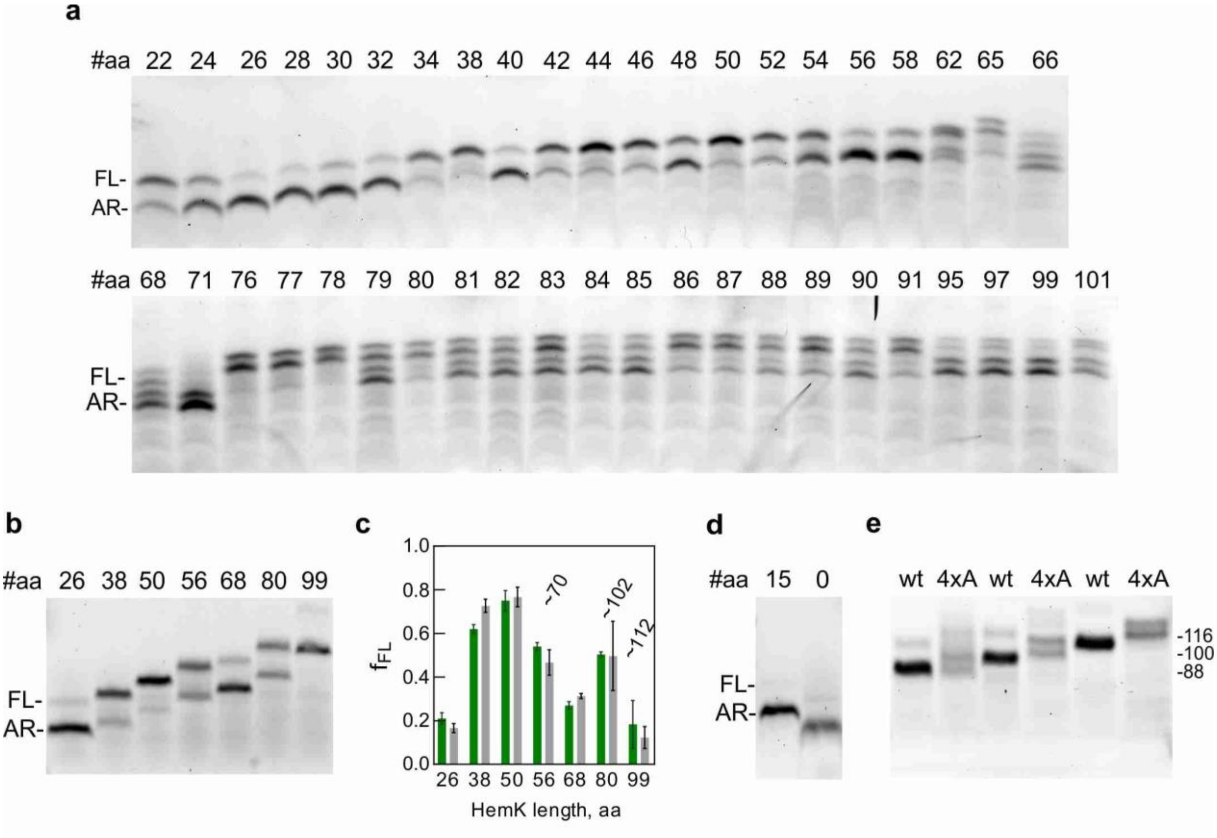

**Supplementary Fig 1** Representative SDS PAGE of FPA for HemK 4xA variant. Abbreviations as in Fig. 1.

a. Representative SDS PAGE of FPA for HemK 4xA variant
b. Representative SDS PAGE of FPA for HemK W6F variant
c. Calculation of fraction of full-length product of HemK W6F (green) compared with wt (grey). Approximate HemK PET-FCS construct lengths indicated on graph. Error bars indicate standard error of mean calculated from three independent biological replicates (N=3)
d. Representative SDS PAGE of the translation product of the mRNA truncated after codon 15 of *HemK* sequence (15 aa) and SecM arrest peptide only control (0 aa)
e. SDS PAGE of HemK wt and 4xA constructs at lengths 116, 100 and 88, HemK length indicated on the right.

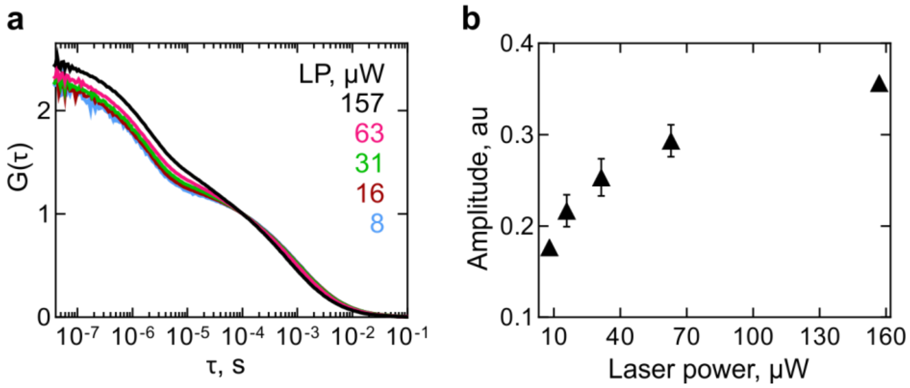

**Supplementary Fig 2** ATTO 655 triplet state in RNC.

a. Autocorrelation curves of HemK102 wt RNC measurements at increasing laser power (LP).

b. Amplitude of the triplet state relaxation time.

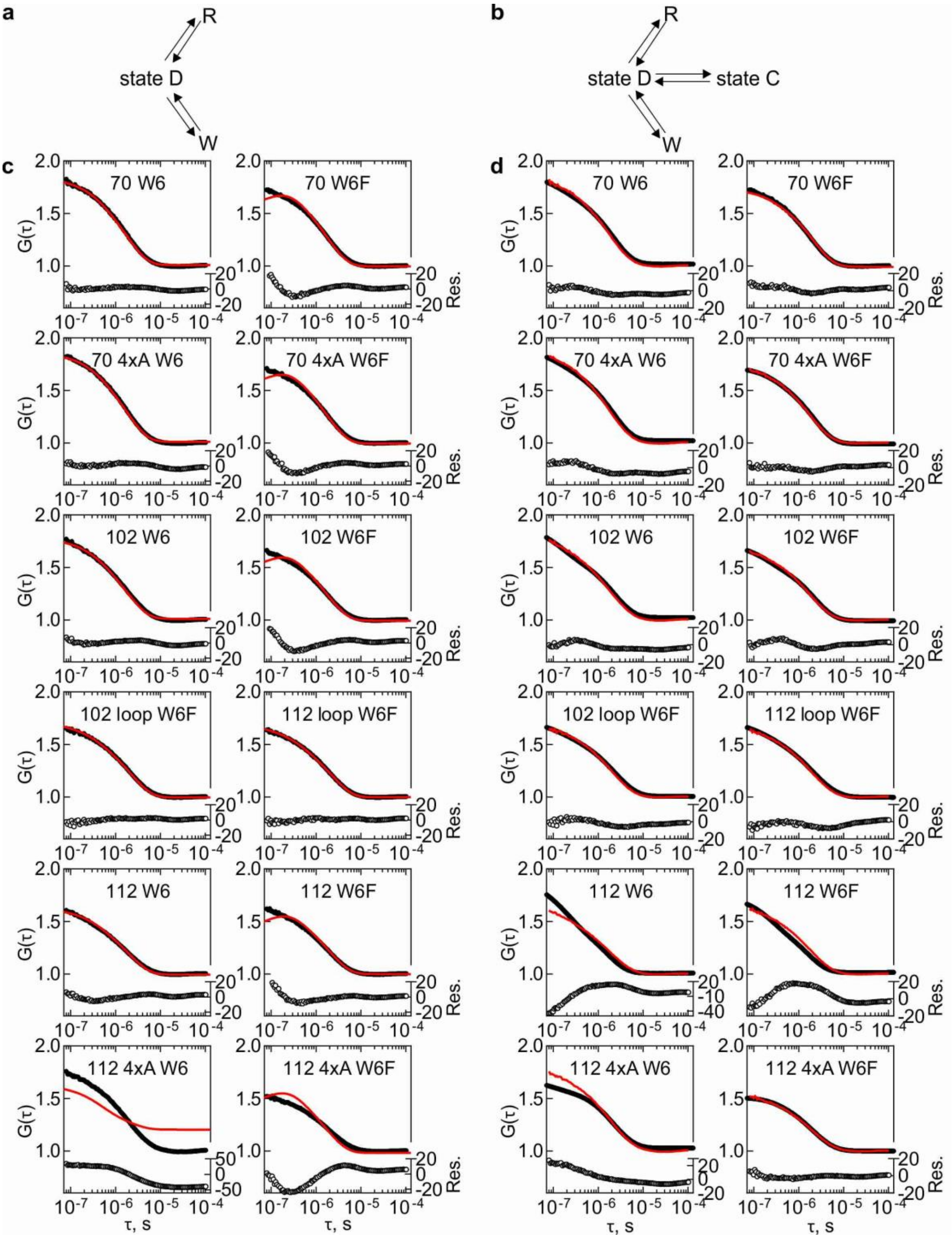

**Supplementary Fig 3** Global fitting of HemK RNC dataset B to kinetic models 2e and 3e.

a. Kinetic Model 2e: nascent chain – state C; Trp-quenched state - W, ribosome-quenched state - R.

b. Kinetic model 3e: nascent chain – state C and state D; W and R same as in a.

c. Results of global fitting of autocorrelation data to kinetic model in 2e, shown in a. Plotted on the left-hand y axis: black – measured data, red – kinetic model simulation curve, On the right-hand y axis in open circles – residual values (Res.); Each graph shows a different RNC construct as indicated.

d. Results of global fitting to kinetic model 3e, shown in b. Graph legend as in c.

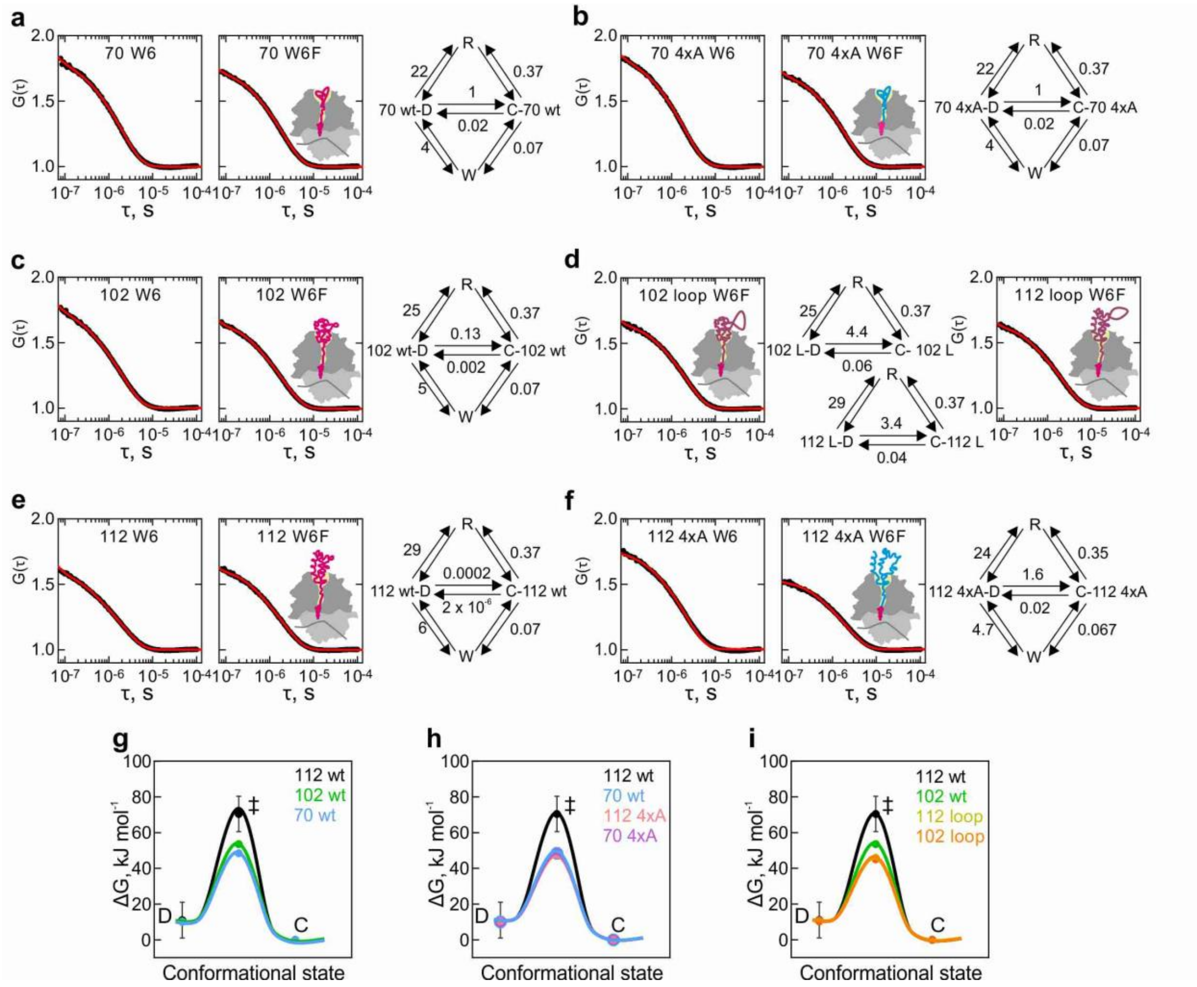

**Supplementary Fig 4** Results of global fitting of HemK dataset B to kinetic model 5-O (right panel). Black – measured data; red – kinetic model simulation curve. C and D are peptide conformations; W – Trp-quenched state; R – ribosome-quenched state. Cartoons illustrate the likely nascent chain position on the ribosome, the wt nascent chains are in magenta, without and with an extra loop; the 4xA variant is shown blue.

a. HemK 70 W6 and W6F.

b. HemK 70 4xA W6 and W6F.

c. HemK 102 W6 and W6F.

d. HemK 102 and 112, both W6F, with loop extensions.

e. HemK 112 4xA W6 and W6F.

f. HemK 112 4xA 4xA W6 and W6F.

Free energy barriers between different chain conformations calculated from model 5e-O rates, values and SEM shown in Supplementary Table 9.

g. HemK wt constructs of increasing length, 70 aa (blue), 102 (green), and 112 (black).

h. HemK wt and 4xA variants, 70 wt (blue) 70 4xA (lilac), 112 (black), and 112 4xA (red).

i. HemK wt and loop variants, 102 wt (green), 112 wt (black), 102 loop (orange), and 112 loop (yellow).

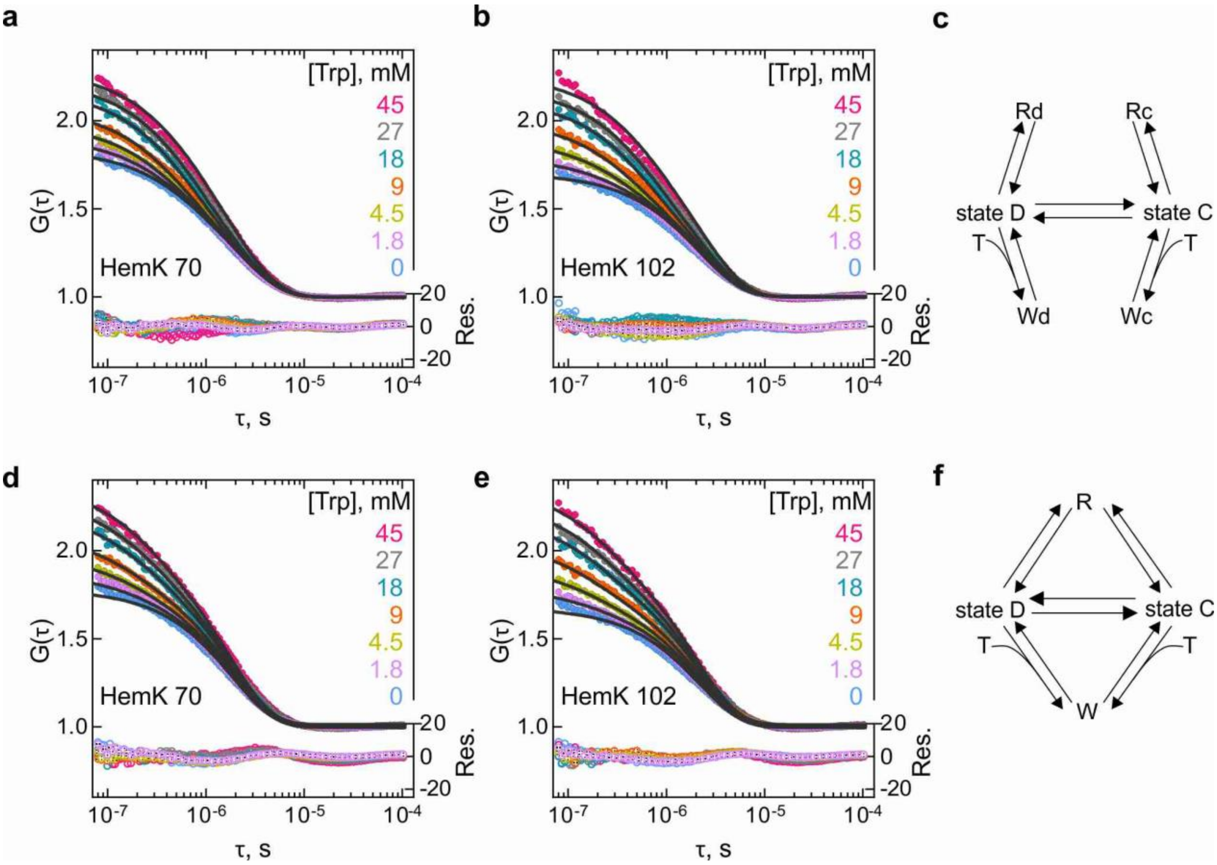

**Supplementary Fig 5** Global fitting of autocorrelation curves for HemK W6F RNCs with increasing free Trp concentration. Free Trp concentration is indicated on the right of each graph.

a. HemK 70 W6F RNC, fitted to model 5e-H (c).
b. HemK 102 W6F RNC, fitted to model 5e-H (c).
c. Global fit kinetic model 5e-H. Nascent chain conformational states – state D and state C; ribosome quenched states Rd and Rc, Tryptophan quenched states Wd and Wc; Tryptophan in solution T.
d. HemK 70 W6F RNC, fitted to model 5e-O (f).
e. HemK 102 W6F RNC, fitted to model 5e-O (f).
f. Global fit kinetic model 5e-O.

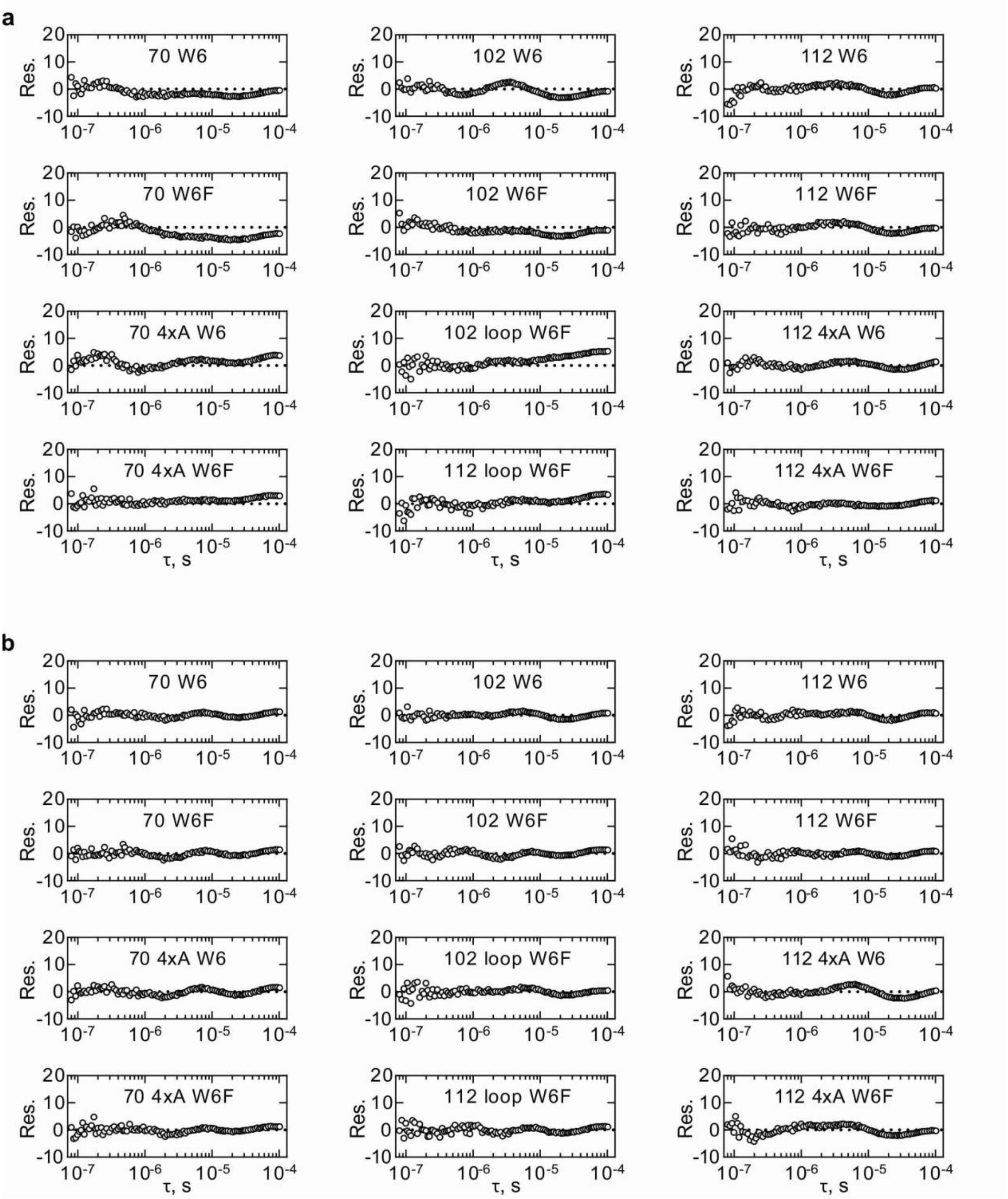

**Supplementary Fig 6** Plots of residuals from the kinetic model 5e fittings of the HemK RNC dataset B.

a. Residuals of fits to model 5e-H.
b. Residuals of fits to model 5e-O.
